# Predictive Functions of H3K27me3 and H4K20me3 in Primate Hippocampal Stem and Progenitor Cells

**DOI:** 10.1101/2021.08.03.454824

**Authors:** Christopher T. Rhodes, Chin-Hsing Annie Lin

**Author notes:** **Corresponding authors:** Chin-Hsing Annie Lin.

## Abstract

Epigenetic regulations play important roles in cell fate determination during neurogenesis, a process by which different types of neurons are generated from neural stem and progenitor cells (NSPCs). Although some epigenetic changes are part of developmental and aging processes, the role of tri-methylation on histone 3 lysine 27 (H3K27me3) and histone 4 lysine 20 (H4K20me3) in primate hippocampal NSPCs remains elusive. This task is best assessed within a context resembling the human brain. As more studies emerge, the baboon represents an excellent model of human central nervous system in addition to their genomic similarity. With a focus on H3K27me3 and H4K20me3, the overarching goal of this work is to reveal their respective epigenetic landscapes in NSPCs of non-human primate baboon hippocampus. We identified putative targets of H3K27me3 and H4K20me3 that suggests a protective mechanism by dual H3K27me3/H4K20me3-mediated repression of specific-lineage gene activation important for differentiation processes while controlling the progression of the cell cycle.

## INTRODUCTION

Within neurogenic niches, a population of neural stem progenitor cells (NSPCs) constantly interacts with niche support and can differentiate stepwise to become mature neurons. Using viral-based lineage tracing and fate-mapping in rodent model ^1,2^, the NSPCs subtypes in the subgranular zone of dentate gyrus (SGZ/DG) were identified, such as GFAP and Nestin as markers for early neural stem/progenitor cells; Doublecortin (DCX) as a marker of neuroblasts. The sequential steps of cell fate transition from GFAP to DCX are regulated by numerous mechanisms including transcription factors, signaling pathways, and epigenetic modifications. Given that the post-translational methylations onto histone tails are influential regulators of cell fate, our previous work had found that H3K27me3 and H4K20me3 are preferentially enriched in undifferentiated hippocampal NSPCs ^3^. H3K27me3 and H4K20me3 are catalyzed independently by the SET-domain superfamily of histone methyltransferases (HMTs) -the enhancer of zeste homolog 2 (EZH2) and the suppressor of variegation 4-20 homolog (Suv4-20h1 and Suv4-20h2), respectively ^4-6^. Both H3K27me3 and H4K20me3 are associated with chromatin compaction and transcriptional repression that can be inheritable to progeny, known as epigenetic inheritance ^6-10^. Yet, their targets in primate hippocampal NSPCs are unclear that require epigenetic mapping using model organisms better mimicking human brain.

While rodent models have been of tremendous use to broad aspects of studies, their brain volumes are much smaller and contain fewer white matter tracts than those of humans. Of the commonly used laboratory primates (baboon, green monkey, macaque, marmoset), the baboon has high similarity to human in terms of brain anatomical structure, size/volume, and brain glucose metabolism ^11-13^. In addition, the baboon has the closest ratio of gray matter versus white matter to human ^11,12^, shares 92% genomic homology and 90% homologous MHC coding loci with humans ^14^, also have all four IgG subtypes as human at 87-90% similarity depending on the subtypes ^15,16^. Further, the overall microvasculature of the baboon brain is highly reflective of that in the human ^11,12,17-19^. These evidences suggest that the baboon provides a viable alternative to the great apes as a model, especially when the United States National Institutes of Health has announced phasing out chimpanzees for research on 2013. Using a non-human primate model baboon (*Papio anubis*), we have shown that the overall architecture and the NSPC expression pattern in the baboon brain phenocopy findings in human brain ^13^. Therefore, we isolated NSPCs from baboon hippocampus to identify putative targets of H3K27me3 and H4K20me3 that infer predictive functions in undifferentiated hippocampal cells.

## RESULTS

To overcome the spatial complexity of SGZ/DG and to ascertain endogenous characteristics of the NSPCs, we developed a technique ^13^ to purify baboon NSPCs from adult hippocampus in a manner that preserves the nature of these undifferentiated cells. Markers representing neural stem cell (GFAP, also known as radial glial-like cells), progenitors (Nestin), and neuroblast (DCX) were applied to NSPCs purification. Briefly, we manually conjugated Dynabeads with antibody recognizing NSPCs-specific markers (GFAP, Nestin, and DCX). We then used the Dynabeads-conjugated antibody to purify cells dissociated from SGZ/DG after microdissection ^13^ (Figure 1). These purified cells without the artifact from cell culture allow us to perform epigenetic mapping of H3K27me3 and H4K20me3 via ChIP-Seq (immunoprecipitation & deep-sequencing) (Figure 1).

**Figure 1:**
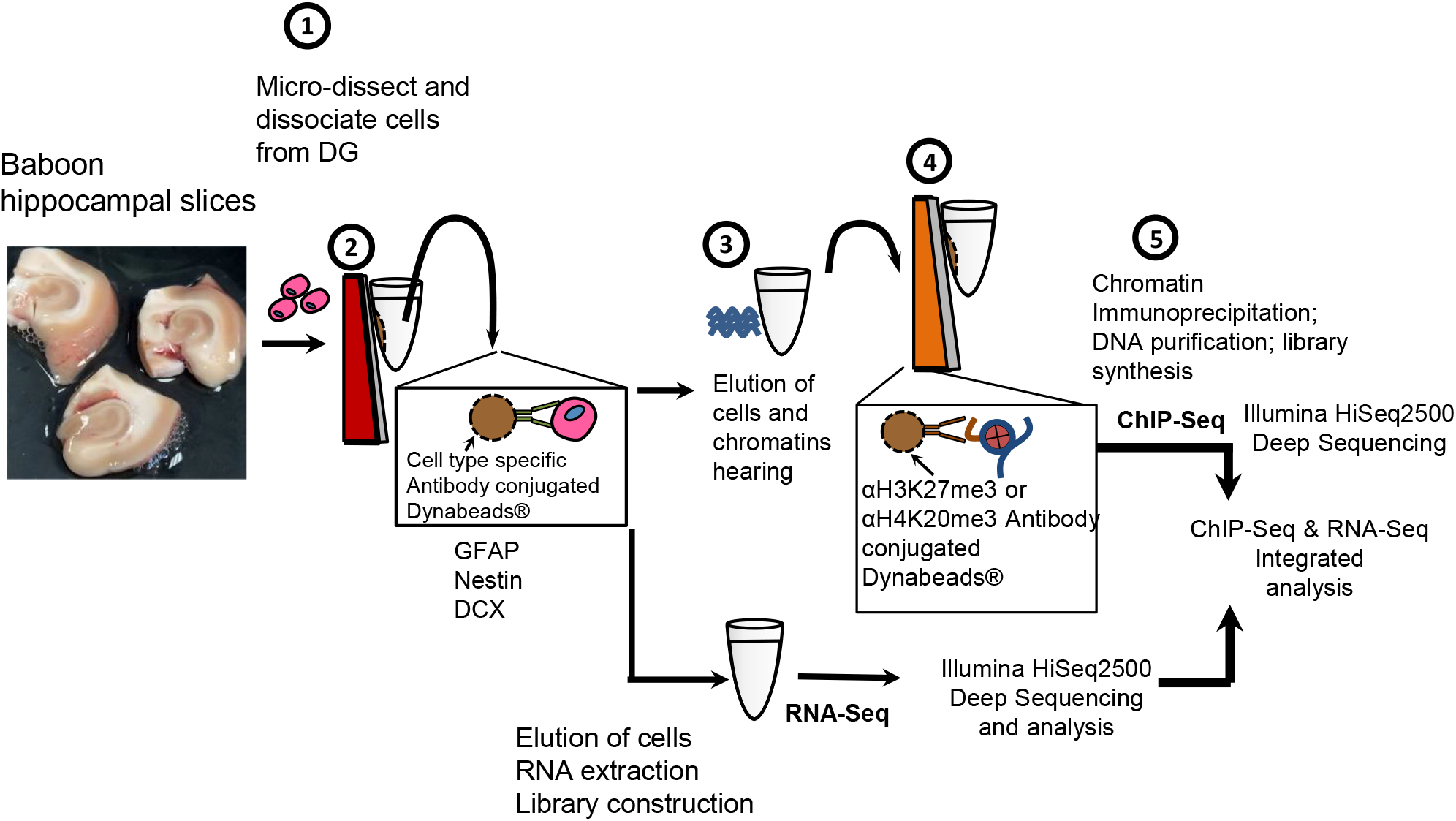
Scheme displays coronal view of baboon hippocampal slices with the dictation of NSPCs isolation prior to the process of ChIP-Seq and RNA-Seq. Briefly, GFAP antibody-Dynabeads were used for NSC purification, Nestin-Dynabeads were used for active progenitor isolation, DCX-Dynabeads were used to purify neuroblasts.

To identify putative targets of H3K27me3 and H4K20me3 in undifferentiated cells residing in the SGZ/DG within the hippocampus, purified NSPCs were subjected to chromatin immunoprecipitation (ChIP) with antibodies recognizing H3K27me3 and H4K20me3, followed by deep-sequencing. H3K27me3 is known to enrich gene promoters and TSSs in bivalent domains ^20^, also is canonically described as broadly enriching gene bodies. As a means of validation, we integrated ChIP-Seq and RNA-Seq data (FDR <0.01) to further confirm putative targets of H3K27me3 and H4K20me3 in the hippocampal NSPCs (Figure 1). Under deep-sequencing level, we obtained approximately 200 million pass-filter reads for ChIP-Seq and 270 million mapped reads (input reads ∼ 300 million) for RNA-Seq. Using Cufflink, the fragments per kilobase of exon per million fragments mapped (FPKM) was applied to RNA-Seq analysis, and FPKM < 1 was determined as non-expressed (repressed) genes before integrating ChIP-Seq and RNA-Seq data. Next, Gene Ontology and Ingenuity Pathway Analysis (IPA) reveal top networks and pathways for genes enriched by H3K27me3 (Figures 2 and 3) and H4K20me3 (Figures 4 and 5). Distinctively, H3K27me3-enriched genes are predominantly involved in the development and differentiation processes (Figures 2 and 3) while H4K20me3 enrichment highlights gene sets in immune response and metabolism (Figures 4 and 5). Furthermore, we uncovered genes co-enriched with both epigenetic repressors that have functions primarily in cell cycle regulation, synapse or glutamate receptor signaling, and chromatin network (Figure 6). This result suggests their collaborative role in hippocampal function.

**Figure 2:**
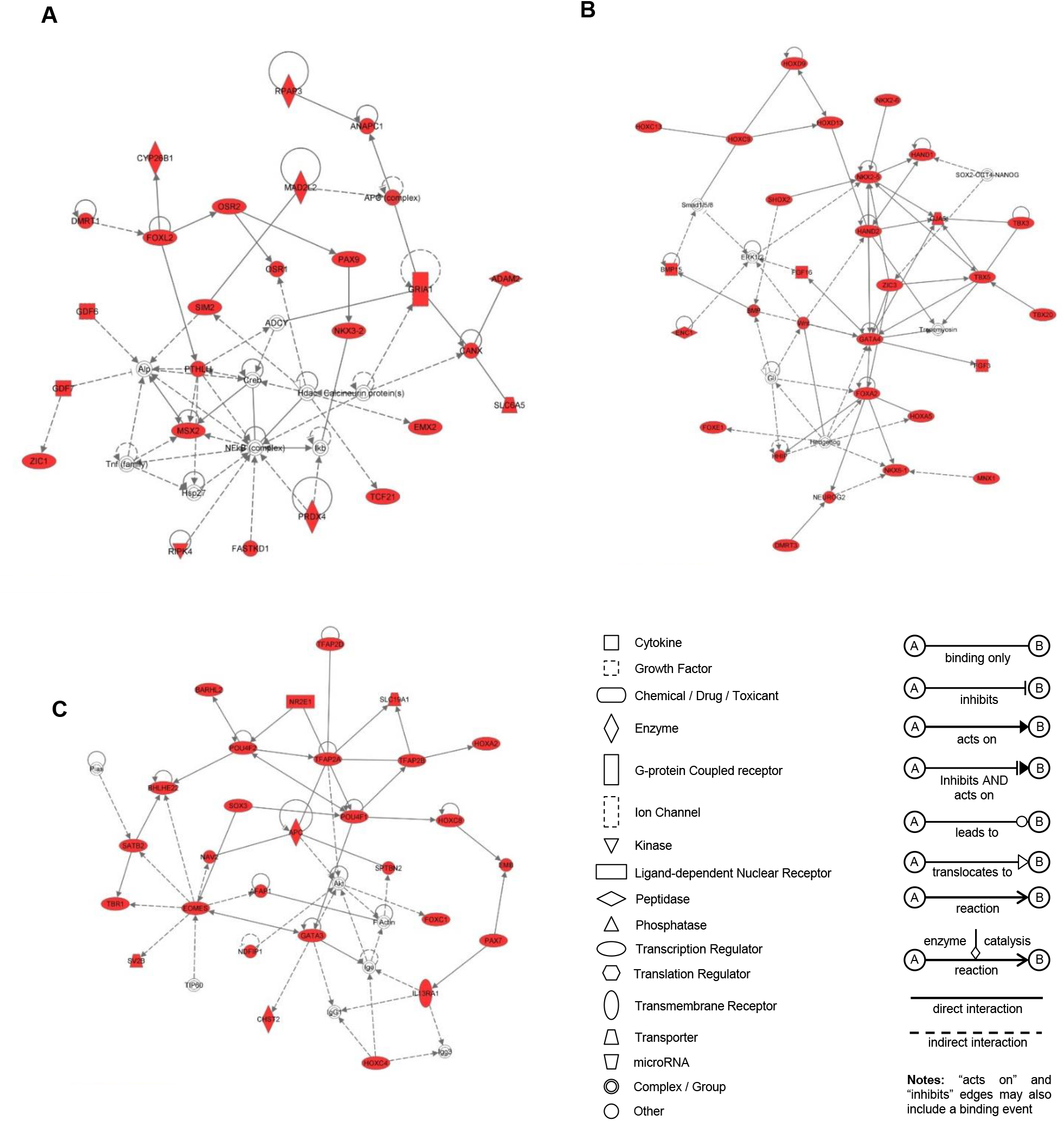
Functional category predicted by IPA for genes enriched with H3K27me3 in hippocampal neural stem and progenitor cells (NSPCs). (A) putative target genes are involved in neurogenesis; (B) nervous system development and function; (C) transcription factors for differentiation and morphogenesis. The shaded focus genes (red highlight) were enriched with H3K27me3. The genes without red highlight were not enriched with H3K27me3, but were involved in the network or pathway. Node shape reflects the role of each gene in the network, i.e. transcription factors, transporters, kinases. The direction and arrowhead shapes of each edge represent different types of interactions.

**Figure 3:**
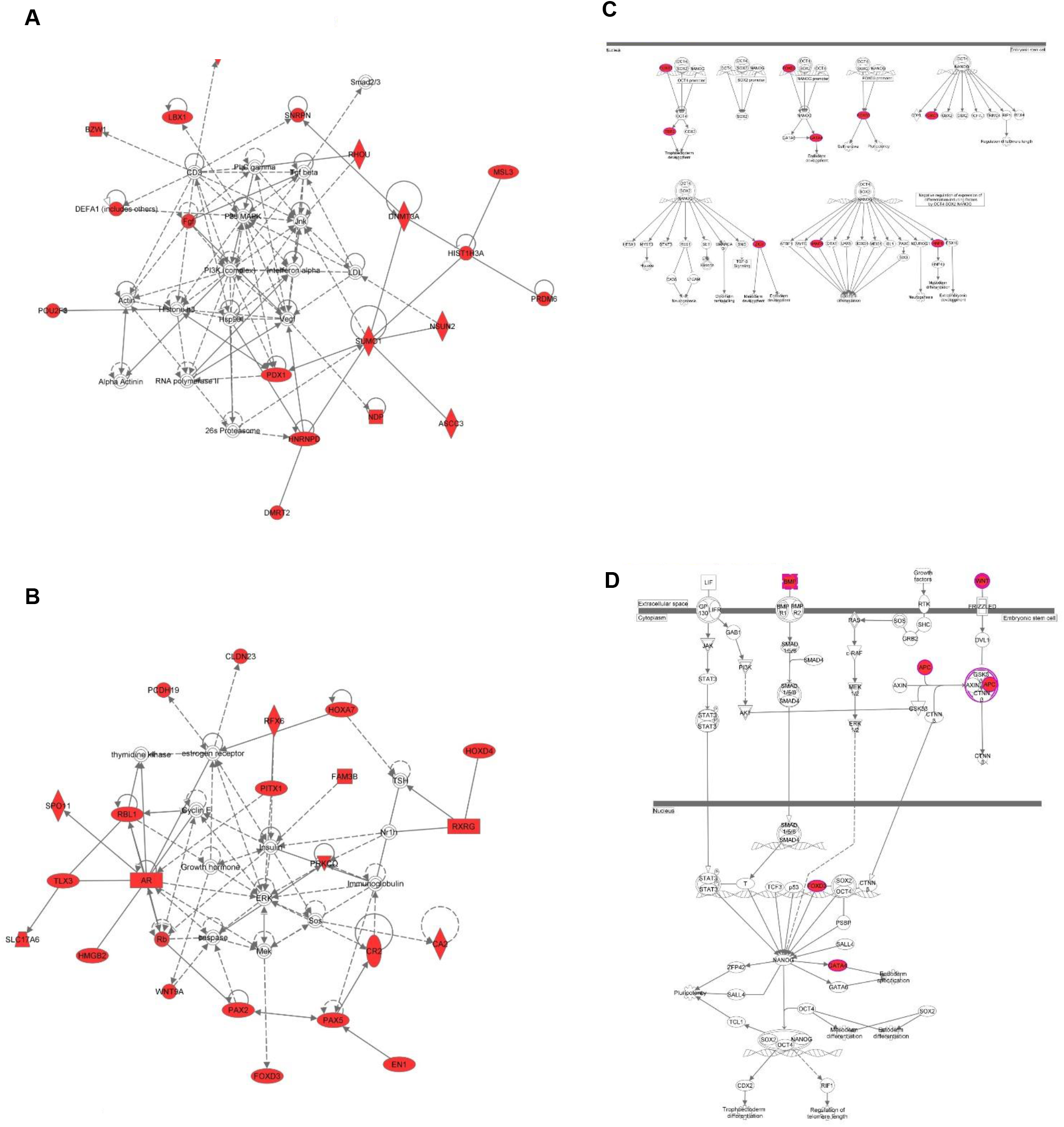
Genes enriched with H3K27me3 in hippocampal NSPCs have functions in pluripotency and development predicted by IPA. Putative targets were associated with (A) cellular maintenance and morphogenesis; (B) cell growth and proliferation; stem cell pluripotency shown in (C) Sox2/Oct4 pathway and (D) Nanog pathway.

**Figure 4:**
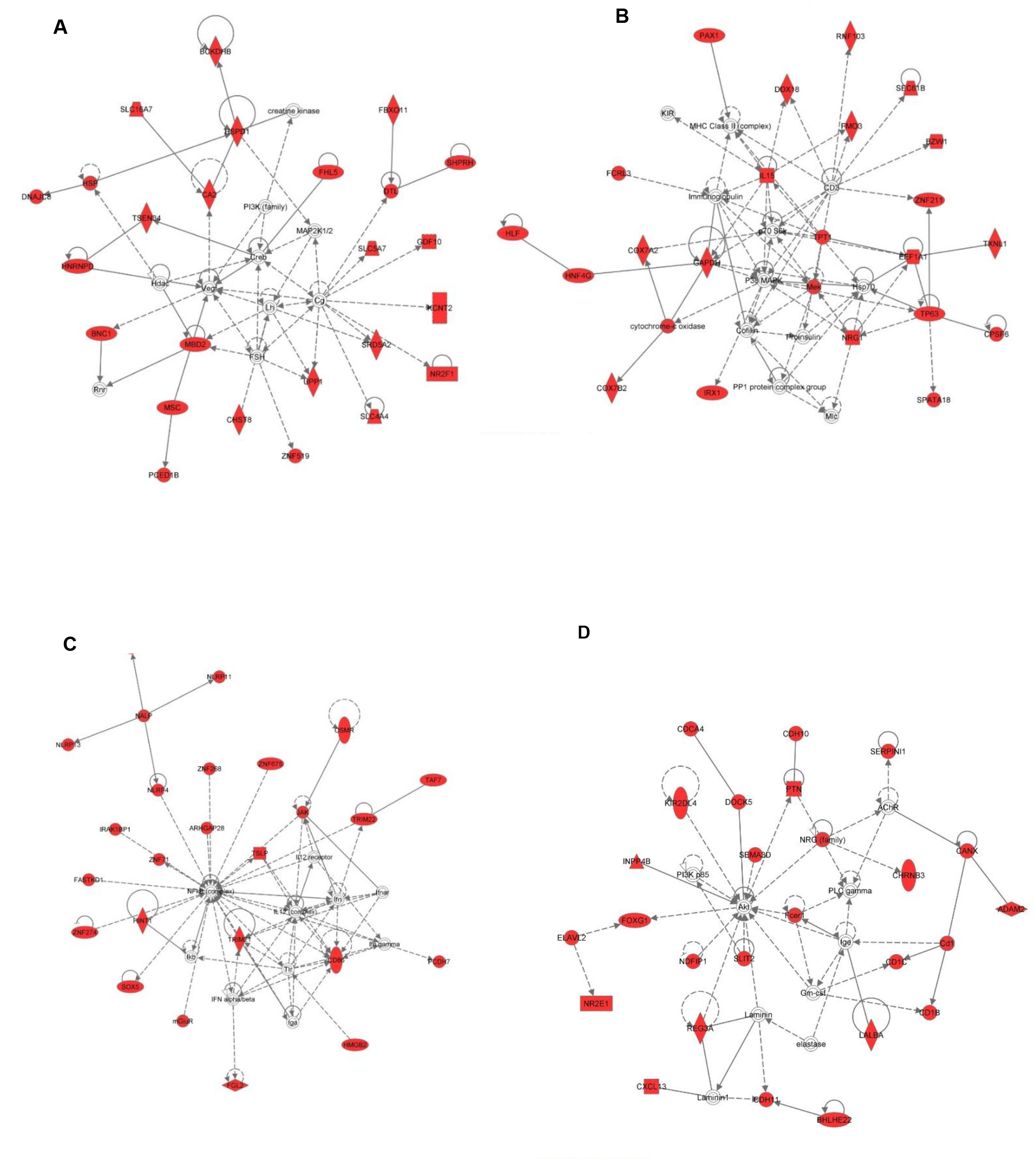
Functional category predicted by IPA for genes enriched with H4K20me3 in hippocampal NSPCs. Putative target genes are involved in (A) nucleic acid metabolism; (B) antigen presenting metabolism; (C) cell death and survival; (D) lipid metabolism.

**Figure 5:**
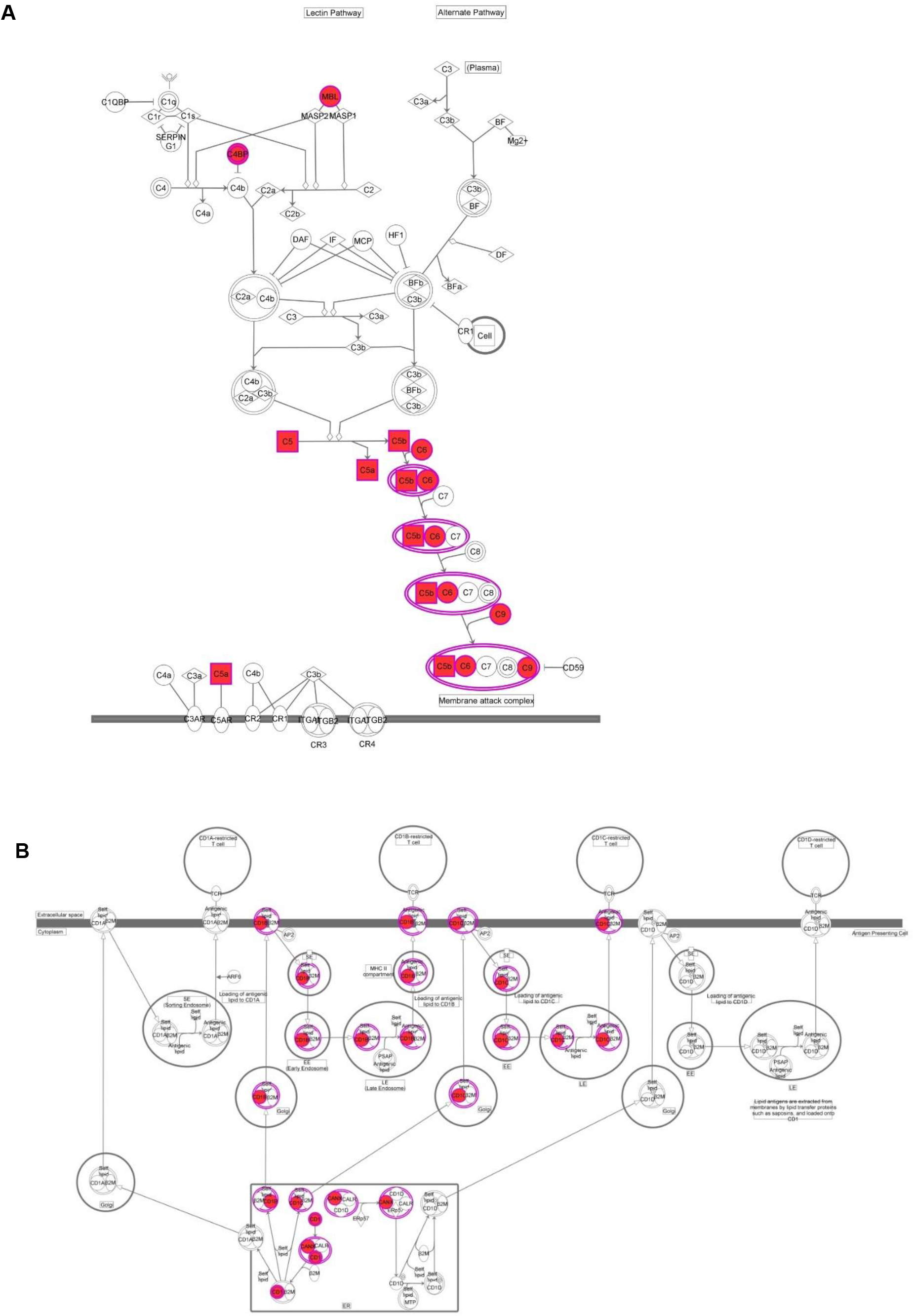
Genes enriched with H4K20me3 in hippocampal NSPCs were associated with immune response. Functional categories predicted by IPA include pathways in (A) complement system; (B) lipid antigen presentation by CD1.

**Figure 6.**
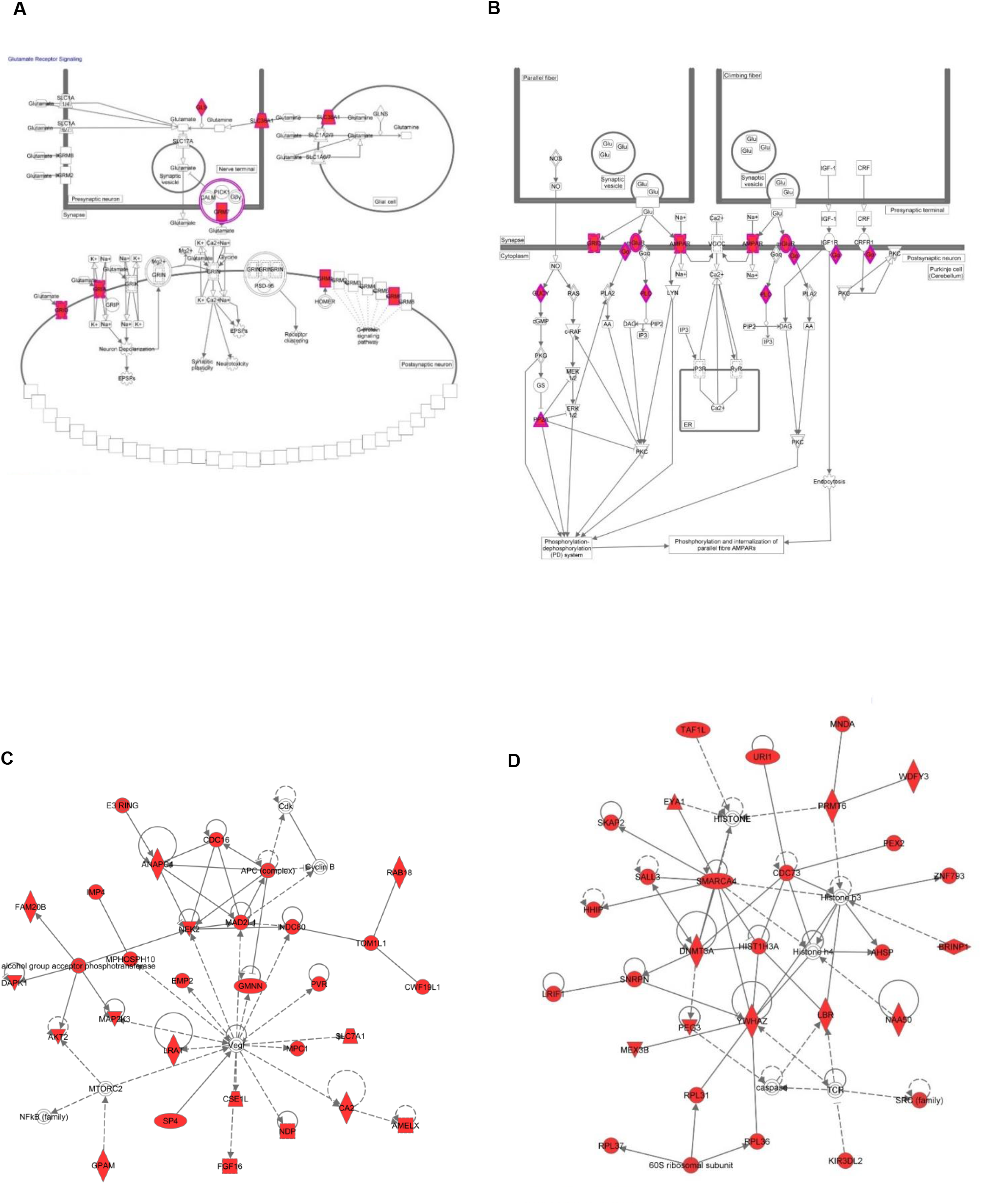
Putative co-target genes of H3K27me3 and H4K20me3 are involved in (A) glutamate receptor signaling for neuron and astrocyte interaction; (B) synaptic long-term repression; (C) cell cycle regulation; (D) chromatin network. The shaded focus genes (red highlight) were enriched with both H3K27me3 and H4K20me3.

## DISCUSSION

Hippocampal stem progenitor cells can potentially generate an inexhaustible supply of neurons important for regeneration. However, all clinical attempts to apply stem cell therapies to humans had failed, despite some success in rodent studies. It is not surprising as rodents have much smaller brain and are limited in their MHC homology to model the human inflammatory response ^21^. Baboons, on the other hand, share 92% genomic sequence homology, better correlate to anatomical structure of human brains, and recapitulate behavioral/cognitive abilities and functional neural circuits as those of humans as well. Importantly, the baboon immune system faithfully represents the human making the baboon a good model to evaluate the risk of immune rejection after cell-based transplantation ^15,16^.

Characterization of the H3K27me3 and H4K20me3 epigenetic landscape is critical to understanding how undifferentiated cells properly divide and differentiate under the control of these two chromatin repressors. Herein, the intellectual merit of this work lies in epigenetic regulations of H3K27me3 and H4K20me3 underlying NSPCs proliferation, cell fate transition, and differentiation in a model system that is much more relevant to humans. As the transition between proliferation and differentiation is frequently associated with changes in chromatin status and gene expression ^22^, our data suggests that the difficulty in creating directed differentiation protocols is attributed to epigenetic repression by EZH2/H3K27me3, which functions to keep stem cells in the undifferentiated state. Because a problem that plagues stem cell therapies is the difficulty in developing precise protocols that yield highly specified neuronal subtypes after differentiation ^23^ and subsequent reinstatement of a functional neuronal network, we hypothesize that manipulating the level of EZH2/H3K27me3 could overcome part of the challenges that hinder cell-based therapies for neurological disorders. Furthermore, the irrefutable role of Suv4-20h/H4K20me3 in immune response implicates its critical aspect of successful cell transplantation therapy. Lastly, both epigenetic regulators may collaboratively act in cell cycle-dependent manner crucial for self-renewal and/or plasticity of hippocampal NSPCs. In summary, given the extensive correlation of brain structure ^11,12^ and genomic similarity between baboon and human ^14^, findings regarding epigenetic regulation in baboon models may hold significant relevance to human. Therefore, our approach will be of considerable interest to those applying cutting edge techniques to phenomena in primate model.

## METHODS

All animal experiments were approved by the guidelines of the Institutional Animal Care and Use Committee of the University of Texas at San Antonio (UTSA) and Texas Biomedical research Institute/Southwest National Primate Research Center (SNPRC) at San Antonio.

### Cell types purification for ChIP-Seq analysis

We manually conjugated antibody specific for cell type markers, such as GFAP, Nestin, or Doublecortin to Dynabeads. We then used the Dynabeads-conjugated antibody to purify cells dissociated from DG. Briefly, cells from fresh dissected baboon DG were immediately dissociated with Accutase, equilibrated in binding buffer containing phosphate-buffered saline (PBS) and 0.01 % NP-40 (or saponin, detergent choice depends upon antibody) before incubating with Dynabeads-conjugated antibody. After elution with high salt and pH-gradient buffer (pH 6.5-7.2), the purified populations were crosslinked in 1.1% formaldehyde before chromatin shearing by Diagenode Bioruptor. The resulting sheared chromatin fragments in a size range between 200 to 500 base pairs were incubated with H3K27me3 (Millipore #07-473,1:1000; Active Motif #39160, 1:500) or H4K20me3 (Abcam, Lot#GR212686-1, 1:1000) antibody-conjugated Protein A Dynabeads (Life Technology Dynabeads protein A) overnight. For normalization, the aliquot of sheared chromatin fragments were incubated with antibody conjugated Protein A Dynabeads using total histone 3- (unmodified H3 antibody, Millipore #05-499; 1:1000) or histone 4- (unmodified H4 antibody, Abcam #ab10158). Subsequently, enriched chromatin fragments were eluted, subjected to de-crosslink, purified for library preparation (Illumina Library Kit), and deep-sequencing with 200 million tags through Illumina HiSeq2500 sequencer.

### Sequence alignment and peak calling

Pass filtered ChIP-Seq reads were aligned to the olive baboon (PapAnu2.0) reference genome maintained by Ensembl (ftp://ftp.ensembl.org/pub/release-78/fasta/papio_anubis/dna/) lacking mitochondrial genome with Bowtie v1.1.1 (http://bowtie-bio.sourceforge.net/index.shtml) aligner, allowing a maximum of 2 mismatches in uniquely mapping reads. Genomic regions enriched with target histone modifications were detected using MACS2 v2.1.0.20140616 (https://github.com/taoliu/MACS/) callpeak function (with a false discovery rate (FDR) < 0.05 for each peak). H3K27me3 enrichment was “mapped read density” after normalization to unmodified H3. Likewise, the H4K20me3 enriched loci was normalized to unmodified H4. Gene associations from peak calling were established in a distance ranked manner by using the closest-features program from the BEDOPS tool set ^24^ (https://code.google.com/p/bedops/).

### RNA-Seq Analysis

Total RNA was extracted from purified baboon DG cells using TRIzol reagent and sequencing libraries were generated with Illumina RNA-Seq library kit. RNA libraries were deep sequenced using paired-end sequencing (2×36 bp, >300 million reads) on an Illumina HiSeq2500 sequencer. Pass filtered reads were aligned to PapAnu2.0 reference genome and transcript annotation using Tophat2 v2.0.13 (https://ccb.jhu.edu/software/tophat/index.shtml). Transcripts were assembled by Cufflinks v2.2.1 (http://cole-trapnell-lab.github.io/cufflinks/) using PapAnu2.0 transcript annotation to guide assembly and read counts reported as FPKM (fragments per kilobase of exon per million fragments mapped).

### GO, network, and pathway analysis

Gene ontology analysis was performed using the DAVID functional analysis tool ^25^. The Bonferroni, Benjamini, and FDR (false discovery rate) were used for multiple test correction. Functional pathway and network analysis of bound loci were performed using Ingenuity Pathway Analysis (IPA) (Ingenuity® Systems, Redwood City, CA, USA), then focus genes were grouped into ontology classes. The Ingenuity Knowledge Base, a repository of biological and chemical interactions, was used as a reference set for pathway and network inference. Over-represented signaling and metabolic canonical pathways in the input data were determined based on two parameters: 1) The ratio of the number of molecules from the focus loci set that map to the pathway divided by the total number of molecules that map to the canonical pathway, and 2) a *p* value calculated by Fisher’s exact test that determines the probability that the association between the focus genes and the canonical pathway is explained by chance alone. Direct or indirect interactions of network or pathway analysis were inferred based on experimentally observed relationships supported by at least one reference from the literature. The score was based on the hypergeometric distribution and calculated with the right-tailed Fisher’s exact test. A higher score indicates a lower probability of finding the observed number of focus molecules in a given network/pathway by chance.

## ACKNOWLEDGMENT

Authors acknowledge Texas Biomedical Research Institute/Southwest National Primate Research Center at San Antonio for baboon tissues and Dr. Jera Pecotte for technical support. This project is supported by the SCORE Grant from the National Institutes of Health (NIH/NIGMS) and TRAC award to CAL.

## Author contributions

CTR performed alignment/peak calls analyses, overlap comparison, GO/network/pathway and gene expression data analyses. CAL designed study and experiments, performed ChIP, and wrote manuscript. All authors reviewed manuscript.

## Additional Information

The authors declare no conflict of interest.

